# Amino acid residues for specific binding to ssDNA facilitate topological loading of bacterial condensin MukB

**DOI:** 10.1101/2023.09.21.558748

**Authors:** Koichiro Akiyama, Koichi Yano, Hironori Niki

**Author notes:** Present Address: [Koichi Yano], Department of Science, University of Rikkyo, Nishi Ikebukuro, Tokyo, 171-8501, Japan.

## Abstract

The bacterial condensin MukB facilitates proper chromosome segregation in *Escherichia coli*. A portion of the MukB proteins localize at a specific chromosome region, binding to DNA in a non-sequence-specific manner. However, it is unclear how MukB localizes at a particular site without sequence specificity. Like other structural maintenance of chromosome (SMC) proteins, MukB topologically loads onto DNA, and It has an intrinsic property of preferential topological loading onto the single-stranded DNA (ssDNA). We consider it crucial for the localization of a specific region. To investigate the property of MukB, we attempted to identify positively charged amino acid residues responsible for ssDNA binding. We created a series of mutated MukB proteins in which a single positively charged amino acid was replaced with a negatively charged one. The results showed that some substitutions located on the inner surface of the MukB head domain impacted ssDNA-binding activity, leading to deficiencies in cell growth and nucleoid segregation. The efficiency of topological loading onto ssDNA was also decreased when the positive charges were replaced with negative ones. These amino acid residues align with and bind to ssDNA when the MukB dimer secures ssDNA within its ring, thereby likely strengthening the ssDNA-binding ability of MukB.

## INTRODUCTION

The compaction of replicated chromosome is crucial for proper chromosome segregation within the confined space of bacterial cells. Nucleoid organization, which is involved in the highly ordered arrangement of chromosomal DNA, plays a critical role in this process. Two key factors contribute to the tight compaction of chromosomal DNA: topoisomerase, which generates DNA supercoiling, and bacterial condensin, which folds newly replicated DNA in a manner that is not yet fully understood (1–4). Bacterial condensin is a member of the structural maintenance of chromosome (SMC) protein family (5–9). Two major types of bacterial condensin have been identified and extensively studied to unravel the molecular mechanisms of DNA compaction: the Smc-ScpAB complex in *Bacillus subtilis* (10–12) and the MukBEF complex in *Escherichia coli* (2, 13).

Smc and MukB are the core units in the Smc-ScpAB and MukBEF complexes, respectively, and are collectively referred to as SMC proteins (5, 7, 8). The N-terminal and the C-terminal parts of the SMC proteins undergo intramolecular assembly, forming the ATPase head domain in an ABC cassette arrangement. In contrast, the middle region forms the hinge domain. The head and the hinge domains are connected by the coiled-coil arm domain. Homodimerization of the MukB protein leads to the formation of a ring structure, a characteristic feature of SMC proteins (5, 6, 14). In the case of the MukB dimer, the flexibility of the arm domain allows the MukB dimer to adopt various shapes (15, 16). The SMC proteins form holo complexes with the kleisin subunits (MukF and ScpA) and the kite subunits (MukE and ScpB) (7, 16–23). In addition to these subunits, MukB interacts with other protein factors such as TopoIV and Acyl-carrier protein (ACP) (14, 24–29).

In the MukBEF complex, the MukB protein possesses the DNA-binding activity. MukE and MukF bind to each other and stably form the MukEF complex, which inhibits the DNA binding of MukB (18, 30). MukB binds to DNA in two modes. One is a canonical protein-DNA interaction mediated by the electrostatic force between the positively charged residue of protein and negatively charged phosphate backbone in polynucleotide. In this mode, MukB binds to single-and double-stranded DNA (ssDNA and dsDNA) by its head domain in a non-sequence-specific manner. Although not all dsDNA-binding proteins possess ssDNA-binding activity, binding to single-or double-stranded DNA is a common feature of some SMC family proteins. However, the biological significance of the ssDNA binding remains elusive. In *B*. *subtilis*, the Smc protein binds to ssDNA through its hinge domain (31. The hinge domain of *B*. *subtilis* Smc can be replaced with that of Rad50, an SMC-like protein in yeast that is involved in DNA repair (32. The ssDNA-binding activity of the *B*. *subtilis* Smc hinge is not essential for the functional cycle of bacterial condensin, if indeed it plays any role at all. In the case of MukB, the significant DNA-binding site is its head domain (19, 33, 34). Several multiple mutations in the MukB head domain impair its dsDNA-binding activity, but it is unclear whether the resulting mutants also have decreased affinity for ssDNA. The other mode is the topological binding, where the protein captures a DNA strand inside its proteinaceous ring (5, 7, 8, 35–37). The topological binding mechanism of the SMC protein family is conserved in bacterial condensin and plays key roles in their properties (38–41).

Understanding the mechanism by which SMC proteins capture chromosomal DNA is essential for elucidating the fundamental and universal principles of their biological functions. The loop extrusion model attempts to explain how SMC proteins fold chromosomal DNA, resulting in an organized loop structure (42–47). Alternatively, it is thought that two SMC proteins, especially when bound topologically to different sites on chromosomes, gather to compact DNA (48. In any case, it is critical to determine where and how SMC proteins topologically load on chromosome sites. However, consensus nucleotide sequences for topological binding have not been found thus far.

MukB is distributed throughout the entire chromosome region except the Ter domain (49, 50), where a MukB elimination system exists due to the MatP protein and *matS* sites (27. However, *in vivo* observations of fluorescent protein-fused MukB showed that a portion of the MukB proteins form clusters at the *oriC*-adjacent region (51, 52). The mechanism by which MukB achieves the localization at a specific site, the *oriC* region, on the chromosome without sequence specificity is still unknown.

On the other hand, *B. subtilis* Smc-ScpAB accumulates on the actively transcribed ribosomal RNA (*rrn*) operon (40, likely through topological-type DNA binding. More than two copies of the *cis*-located *rrn* gene operon are necessary to segregate the sister chromosomes properly (40. By assembling at independent *rrn* regions, Smc-ScpAB complexes contribute to the organization of the *oriC* region. The *rrn* region forms an R-loop, a DNA-RNA hybrid, during transcription, resulting in the relatively stable presence of an ssDNA region (53. Many bacteria, including *E. coli* and *B. subtilis,* possess multiple *rrn* operons near the *oriC* region. One plausible scenario to explain the localization of bacterial condensin to the *rrn* region could involve leveraging the property of ssDNA-specific topological binding, as demonstrated with MukB.

MukB prefers ssDNA to dsDNA as a substrate for topological binding (39. Moreover, other SMC proteins have also been shown to have the potential to capture ssDNA. In yeast, condensin exhibits ssDNA-binding activity and demonstrates an in vitro reaction to promote effectively renaturation of complementary ssDNA (54. The yeast condensin accumulates on rDNA (55, 56). On the other hand, yeast cohesin captures DNA twice sequentially, enclosing two DNA molecules in the ring (57. While both ssDNA and dsDNA are effectively captured at the first topological-binding reaction, only ssDNA can serve as the substrate for the second capturing reaction. These observations strongly suggest that the SMC protein can distinguish ssDNA from dsDNA through an unknown mechanism. In this study, we identify the amino acid residues contributing to MukB’s preference for ssDNA and propose a working hypothesis for the molecular mechanism underlying the preferential establishment of topological loading on ssDNA rather than dsDNA.

## MATERIAL AND METHODS

### Bacterial strains and plasmids

Bacterial strains and plasmids used in this study are listed in Table S1 and S2. LB-broth (10 g/L Bacto tryptone, 5 g/L yeast extract, 5 g/L NaCl: pH adjusted to 7.5 by using NaOH) was used for bacterial culture. Kanamycin (25 µg/mL) was added to medium if necessary. The Δ*mukB*::*cat* mutation was introduced into the MC1061 strain by the λRED recombination method (58, resulting in YAN4081.

To obtain a mutated gene with a single amino acid substitution, the *mukB* gene was cloned into pET28 and expressed as an N-terminus histidine-tagged product, resulting in pHis6-MUKB. The introduction of a single amino acid substitution in His_6_-MukB was achieved through two types of PCR-mediated site-directed mutagenesis. The plasmid DNA of pHis6-MUKB was amplified by PCR using a pair of complementary primers that included the desired mutations. After removing the template DNA of pHis6-MUKB through *Dpn*I treatment, the PCR product was transformed into DH5α cells. Instead of using conventional site-directed mutagenesis, the *in vivo E*. *coli* cloning method (iVEC) was employed (59. DNA fragments having homologous overlap regions at both the 5’ and 3’ ends were amplified through PCR reaction with pHis6-MUKB as a template DNA and a pair of primers, one of which contained the target mutation. The PCR product was then transformed into the iVEC strain using a method involving PEG 8000, as described in Nozaki and Niki (59. The circularization of the PCR fragment by enzymes in the iVEC strain resulted in establishment of a plasmid. For both methods, the established plasmid was extracted from the cell, and the entire coding region of *his_6_*-*mukB* was sequenced.

### Cell viability

Temperature-sensitive growth was determined by assessing cell viability at a specific temperature at which colonies formed on an agar plate. The cells were grown in L-broth at 25°C until the early log phase. A portion of the culture was then removed and washed with saline. The cells were resuspended in saline and serially diluted. Three μL aliquots of each dilution of the cells suspension were then spotted onto multiple L-plates, and respectively incubated at 25°C and 37°C.

### Microscopy

Cells were grown in L-broth at 25°C until the early log phase. An aliquot of the cells was diluted with fresh L-broth followed by further shaking for 2.5 h at 37°C. An additional aliquot of the cells was removed and washed with saline. The cells were resuspended in saline and spread on a poly-lysine-coated slide glass. After air-drying for 7 min, the cells were treated with 80% methanol for 7 min, followed by a wash with ultrapure water and subsequent air-drying. The cells mounted with DAPI containing solution were analyzed using a microscope (ZEISS). Images were processed by using AxioVision or ZEN.

### Purification of His-tagged MukB

The His-tagged MukB protein was prepared essentially as described previously (39.

### Electromobility shift assay (EMSA)

A binding reaction was performed using 10 μL of a reaction mixture comprising a specific quantity of purified His_6_-MukB protein and either 5 fmol of circular single-stranded (css) DNA or covalently closed circular (ccc) DNA from pUC119. The reaction mixture contained a buffer with 25 mM HEPES-KOH (pH 7.6), 100 mM KCl, and 1 mM DTT. After incubating the mixture at 37°C for 20 min, loading dye was added. Subsequently, the reaction mixture was subjected to electrophoresis through a 0.7% agarose gel in Tris-borate EDTA buffer at 100 V for 30 min at 4°C. Prior to loading the reaction mixture, the gel was pre-run at 100 V for 60 min at 4°C. The DNA gel was stained with SYBRGreen II and DNA bands were detected using a LuminoGraph (GE) system. The intensity of DNA bands was measured using Fiji software, which can be found at this website: https://imagej.net/software/fiji/.

### Purification of DNA substrates for EMSA and topological-binding assay

The DNA substrates were prepared following a procedure similar to that described previously (39. Circular single-stranded (css) DNA of pUC119 was generated using M13 phage. *E*. *coli* MV1184 cells containing pUC119 were grown overnight in 3 mL of 2x YT medium (Bacto tryptone 16 g/L, yeast extract 10 g/L, NaCl 10 g/L, pH 7.6) with 0.25% glucose at 37°C. Then, 1.5 mL of the overnight culture was mixed with the lysate of the M13KO7 helper phage and incubated at 37°C for 1 h for infection. The mixture was then transferred into 150 mL of 2x YT medium with 0.25% glucose and incubated at 37°C for 1.5 h. Kanamycin (final concentration 50 μg/mL) was added, and the incubation was continued until full growth was achieved. The culture was centrifuged, and the supernatant was collected. Phage particles in the supernatant were precipitated by ultracentrifugation at 100,000 rpm for 30 min using a TLS-110 rotor (Optima TLX; Beckman Coulter Inc.). The resulting pellet was suspended in 800 μL of Tris-EDTA solution (pH 8.0). The solution was treated with 5 μg/mL RNaseA and 70 unit/ml DNase I (Takara) for 1 h at 37°C. Circular single-stranded DNA of pUC119 was isolated by phenol and phenol/chloroform extraction followed by ethanol precipitation. Covalently closed circular (ccc) DNA was prepared by using a DNA extraction kit (Promega). The obtained cccDNA was purified by CsCl density gradient ultracentrifugation.

### Topological-binding assay

A topological-binding reaction was performed using 10 μL of a reaction mixture comprising 3.6 pmol of His_6_-MukB and 100 ng of css pUC119 DNA. The reaction mixture contained a buffer with 25 mM HEPES-KOH (pH 7.6), 25 mM KCl, 1 mM DTT, and 1 mM MgCl_2_. After the mixture was incubated at 37°C for 15 min, it was chilled on ice and then the reaction was quenched by adding 500 μL of CP buffer (25 mM HEPES-KOH (pH 7.6), 500 mM KCl, 1 mM DTT, 1 mM MgCl_2_, 5% glycerol, 0.35% Triton X-100, and 5 mM imidazole). Then 10 μL of His-Tag beads (Dynabeads; Thermo Fischer Scientific) equilibrated with CP buffer were added and the mixture was gently rotated at 4°C for 30 min. The beads were collected on a magnetic stand and washed with 1 mL of CW1 buffer (25 mM HEPES-KOH (pH 7.6) 750 mM KCl, 1 mM DTT, 1 mM MgCl_2_, 0.35% Triton X-100, and 5 mM imidazole) twice followed by washing with 1 mL of CW2 buffer (25 mM HEPES-KOH; pH 7.6, 100 mM KCl, 1 mM DTT, 1 mM MgCl_2_, 0.1% Triton X-100, 5 mM imidazole). The remnant buffer was completely removed. Then the beads were resuspended in 12 μL of elution buffer (25 mM HEPES-KOH (pH 7.6), 100 mM KCl, 1 mM DTT, 1 mM MgCl_2_, and 500 mM imidazole) and incubated at 37°C for 5 min. To a 10 μL aliquot of the supernatant were then added 2 μL of loading dye (NEB, #B7021S) and 1 μL of 2% sodium *N*-dodecanoylsarcosinate. The samples were electrophoresed (100 V, 20 min, room temperature) and DNAs stained by SYBRGreen II were detected by LuminoGraph (GE).

## RESULTS

### A single amino acid substitution at negatively charged residues on the inside surface of the MukB head domain

MukB preferentially entraps ssDNA rather than dsDNA (39. The topological entrapment of DNA by the MukB ring implies that ssDNA interacts with amino acid residues on the inner surface of the closed proteinaceous ring. Generally, a phosphate residue of ssDNA interacts with a positively charged amino acid residue when a protein specifically binds to ssDNA. The inside surface of the closed MukB dimer carries a predominantly positive charge, while the outer surface is negatively charged (**Figure 1B**). Hence, we focused our attention on the positively charged residues, Lys (K) and Arg (R), on the inner surface of the MukB, as a key element for ssDNA binding. We mainly based our estimation of candidate amino acid residues on the crystal structure of the MukB protein encoded in *Haemophilus ducreyi*, given the high similarity in amino acid sequence between *H. ducreyi* MukB and *E*. *coli* MukB: 77% for the entire region and 89% for the head domain (**Figure 1A**). Within the head domain of the MukB ring, the region ranging from R61 to K119 appears to be mainly localized at the inner surface of the closed proteinaceous ring, which is where the amino acid residues having the potential to interact with ssDNA lie. This region contains 12 positively charged amino acids that lie either on or in close proximity to the inner surface (**Figure 1A and B**). These positively charged residues were found to be well conserved among MukB homologs (**Figure 1A**).

**Figure 1.**
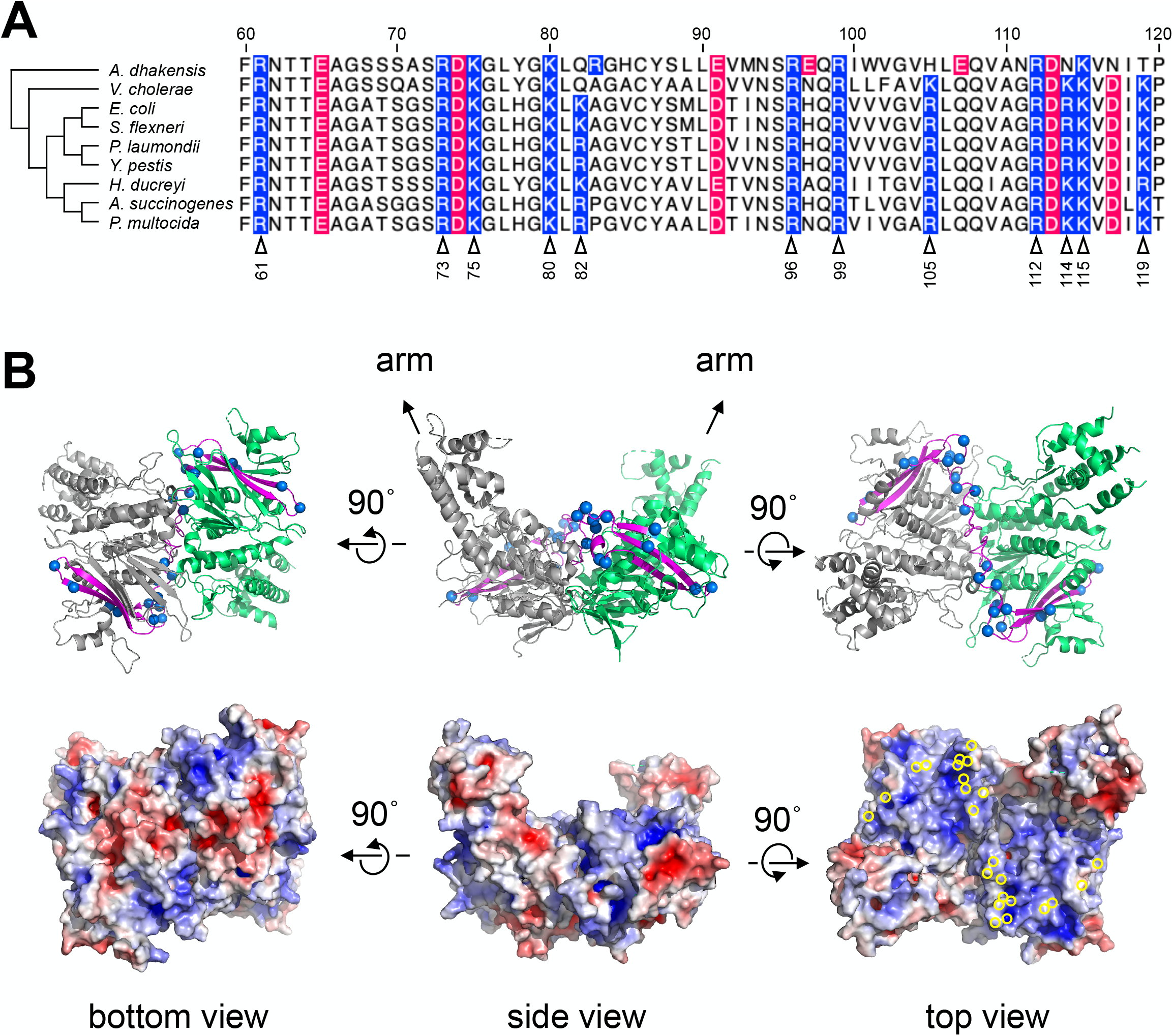
Conserved positively charged residues and crystal structure in the head domain of MukB. (**A**) The phylogenetic tree and the amino acid sequence alignment of MukB homologs indicate nine bacteria closely related to *Escherichia coli*: *Actinobacillus succinogenes*, *Vibrio cholera*, *Escherichia coli*, *Shigella flexneri*, *Photorhabdus laumondii*, *Yersinia pestis, Haemophilus ducreyi*, *Aeromonas dhakensis*, and *Pasteurella multocida*. The alignment, along with the phylogenetic tree, was generated by Clustal Omega (https://www.ebi.ac.uk/Tools/msa/clustalo/). The residue numbers of *E. coli* MukB are indicated at the top of the panel. In the alignment, positively charged residues are highlighted in blue, and negatively charged residues in red. The 12 residues indicated by the open triangles on the bottom with the residue number were substituted in this experiment. (**B**) The crystal structure of the head domain of the *Haemophilus ducreyi* MukB dimer (PDB-ID: 3EUK) is depicted using ribbon diagrams in the top panels, while surfaces with electronic charge density are shown in the bottom panels. These crystal structure models were generated using PyMOL (https://pymol.org/2/). In the top panels, each monomer of the head-engaged dimer is represented in green and grey, respectively. Arrows show the regions connected to the arm. The structure between R61 and K119 is displayed in magenta, and the 12 positively charged residues in panel A are indicated as blue spheres in the top panels and as yellow open circles in the bottom right panel.

We introduced amino acid substitution mutations into the head domain of MukB to determine which of these positively charged amino acid residues on the inner surface of the MukB ring is involved in the specific recognition of ssDNA. A plasmid containing the *mukB* gene with an amino acid substitution mutation was constructed. The plasmid expressed the N-terminal His-tagged gene product under the T7 promoter. Twelve positively charged amino acids were substituted for glutamic acid (E), which is negatively charged, or glutamine (Q), which is neutrally charged. To assess the biological significance of the amino acid substitution in the MukB protein, we examined whether the mutations seriously affect cell growth and chromosome segregation (**Table 1 and Figures 2, 3 and S1**).

**Figure 2.**
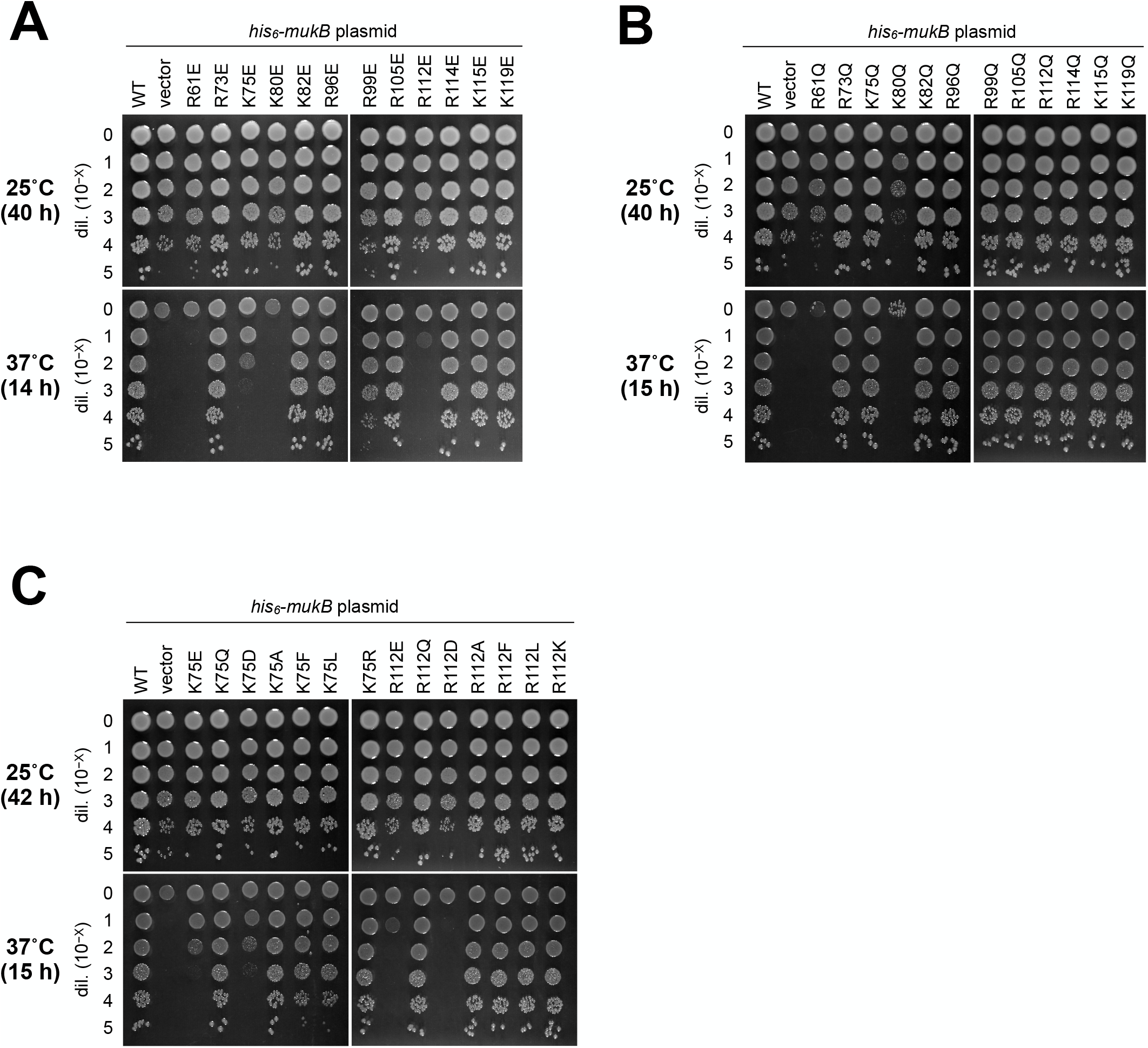
Colony formation test of the MukB head domain mutant-expressing cells. (**A**-**C**) YAN4081 (Δ*mukB*::*cat*) harboring the plasmid encoding the indicated His_6_-MukB mutant was cultured in L-broth at 25°C. Cells were serially 10-fold diluted with saline and spotted on L-plates followed by incubation at 25°C or 37°C for indicated time.

**Table 1.** Properties of the MukB head domain mutants. *a)* The frequency of anucleate cells in relation to the total cell count was determined from photographs as shown in **Figure 3** and **Supplementary Figure 1**. The total cell count was at least 162 and usually exceeded 300 cells. *b)* Defects in nucleoid segregation were assessed from the histogram showing the frequencies of anucleate cells (**Supplementally Figure 2**). In normal nucleoid segregation (+), the percentage is less than 3% at 25°C and 37°C. *c)* The cell shape was clarified from the level of cell elongation; cell size in the wild-type strain (+), a few folded cell lengths of WT (L), extremely elongated cells as filamentous (F). *d)* Cell viability was assessed from the results in **Figure 2**.

| His<sub>6</sub>-MukB | Anucleate cell production rate (%)<sup>a</sup> |  | Nucleoid segregation<sup>b</sup> | Cell shape<sup>c</sup> | Cell viability<sup>d</sup> |
| --- | --- | --- | --- | --- | --- |
|  | 25°C | 37°C | 25°C/37°C | 25°C/37°C | 37°C |
| WT | 0.2 | 0.3 > | +/+ | +/+ | + |
| vector | 6.5 | 8.1 | -/- | L/F | - |
| R61E | 9.2 | 9.6 | -/- | L/F | - |
| R61Q | 10.4 | 13.7 | -/- | L/F | - |
| R73E | 1.4 | 1.3 | +/+ | +/L | + |
| R73Q | 0.5 | 0.5 | +/+ | +/L | + |
| K75E | 3.4 | 5.0 | -/- | L/F | - |
| K75Q | 0.5 | 1.0 | +/+ | L/L | + |
| K75D | 6.2 | 12.7 | -/- | L/F | - |
| K75A | 0.5 | 1.1 | +/+ | +/L | + |
| K75F | 2.5 | 7.7 | +/- | L/L~F | + |
| K75L | 2.4 | 9.3 | +/- | L/L~F | + |
| K75R | 1.2 | 2.1 | +/+ | L/L | + |
| K80E | 4.8 | 14.8 | -/- | L/F | - |
| K80Q | 13.6 | 16.3 | -/- | L/F | - |
| R96E | 0.3 | 1.3 | +/+ | +/L | + |
| R96Q | 1.2 | 0.8 | +/+ | +/L | + |
| R99E | 8.4 | 9.7 | -/- | L/F | + |
| R99Q | 1.3 | 1.3 | +/+ | +/L | + |
| R105E | 1.4 | 1.6 | +/+ | +/L | + |
| R105Q | 0.2 | 0.4 | +/+ | +/L | + |
| R112E | 8.2 | 5.1 | -/- | L/F | - |
| R112Q | 1.7 | 1.5 | +/+ | L/L | + |
| R112D | 7.4 | 8.8 | -/- | L/F | - |
| R112A | 0.5 | 0.2 > | +/+ | L/L | + |
| R112F | 0.8 | 1.0 | +/+ | L/L | + |
| R112L | 1.0 | 0.3 | +/+ | +/L | + |
| R112K | 0.3 | 1.2 | +/+ | +/L | + |
| R114E | 0.7 | 1.2 | +/+ | +/L | + |
| R114Q | 0.2 | 0.5 | +/+ | +/L | + |
| K115E | 1.1 | 0.7 | +/+ | L/L | + |
| K115Q | 0.4 | 0.3 | +/+ | L/L | + |
| K119E | 0.2 > | 0.2 > | +/+ | L/L | + |
| K119Q | 0.7 | 0.2 > | +/+ | L/L | + |

a) Frequencies of anucleate cells in relation to the total cell count were determined from photo images, such as Figure 3 and Figure S1. The total cell count is at least 162 and usually exceeds 300 cells.
b) Defects in nucleoid segregation were assessed from the histogram showing frequencies of anucleate cells (**Supplementally Figure 1**). In normal nucleoid segregation (+), the percentage is less than 3 % at 25°C and 37°C.
c) Cell shape was clarified from the level of cell elongation; cell size in the wild type strain (+), a few folded cell lengths of WT (L), extremely elongated cells as filamentous (F).
d) Cell viability was assessed from the results in **Figure 2**.

### Temperature-sensitive growth of the amino acid substitution mutants

The plasmids encoding the mutated MukB protein with a single amino acid substitution were introduced into the *mukB*-deletion mutant to assess the biological functions of the mutated MukB protein. The *mukB*-deletion mutant shows two representative phenotypes: temperature-sensitive growth and aberrant chromosome separation. Initially, we examined whether the amino acid-substituted mutants could suppress the temperature-sensitive growth of the *mukB*-deletion mutant cells. Cells harboring a plasmid encoding the N-terminal *his*-tagged *mukB* gene under the T7 promoter were incubated at the permissive (25°C) or the restrictive temperature (37°C). As a result, the plasmid encoding the *his_6_-mukB^wt^* enabled the deletion mutant to grow at 37°C, even without addition of an inducer to the agar medium to activate transcription from the T7 promoter (**Figure 2A**). The leaked expression of the His-tagged MukB protein was sufficient for growth of the *mukB*-deletion mutant at 37°C. Similarly, we tested a total of 24 amino acid substitution mutants for their temperature-sensitive growth at 37°C (**Figures 2A and B**).

All the substitution mutants grew at 25°C, but 6 mutants failed to grow at 37°C. Specifically, the amino acid substitution mutations R61E, R61Q, K75E, K80E, K80Q, and R112E were unable to compensate for the temperature-sensitive growth of the *mukB*-deletion mutation (**Figures 2A and B**). Neither the substitution E nor the substitution Q for R61 and K80 could restore the temperature-sensitive growth. These findings suggest that the arginine residue at R61 and the lysine residue at K80 are absolutely necessary for the biological function of the MukB protein. Previously reported structural information suggests that these residues directly participate in and are essential for the ABC transporter-type ATPase activity (19, 60). Therefore, the presence of positively charged residues at these positions is required for the proper functioning of the MukB protein.

On the other hand, the amino acid-substitution mutations K75Q and R112Q were able to compensate for the temperature-sensitive growth, while the substitution mutations K75E and R112E could not (**Figures 2A and B**). To investigate whether the defects observed in the K75E-and R112E-substitution mutants were specifically due to the inversion of charge from positive to negative, we additionally constructed amino acid-substitution mutants of K75 and R112 using various residues: the negatively charged residue Asp (D), the non-charged residues Ala (A), Phe (F), and Leu (L), and the positively charged residues Arg (R) and Lys (K). The results revealed that the amino acid-substitution mutations K75D and R112D, which change the positive charge to a negative charge, failed to suppress the growth defect at 37°C (**Figure 2C**). Conversely, when the positive charge was maintained, as in the case of the amino acid substitutions K75R and R112K, the complementation activity of the MukB protein for temperature-sensitive growth remained unaffected. Additionally, substitutions with non-charged residues did not impact the activity. Therefore, it is concluded that only the presence of negatively charged amino acid residues at positions K75 and R112 had a detrimental effect on the biological function of the MukB protein with respect to temperature-sensitive growth.

### The defect in chromosome separation by the amino acid substitution mutants

We next examined whether the amino acid substitution mutations could reduce the production of anucleate cells in the *mukB* gene-deletion strain. The *mukB*-deletion mutant displayed a high frequency of anucleate cell production, as well as cells with aberrant nucleoids, even when cultured at the permissive temperature. The frequency of anucleate cell production increased and the cells became filamentous when cultured at the non-permissive temperature of 37°C. However, these phenotypes of the *mukB* gene deletion were effectively suppressed by the plasmid encoding the N-terminal *his*-tagged *mukB* gene (**Figure 3**). When the cells carried the vector plasmid, the frequency of anucleate cell production was 8.1% at 25°C (**Figure 3**, **Table 1**). However, the amino acid-substitution mutations R61E, R61Q, K75E, K80E, K80Q, R99E and R112E were unable to suppress the production of anucleate cells (**Figure 3 and Figure S2**). These results were consistent with the findings from the temperature-sensitive growth analysis, except for the amino acid-substitution mutation R99E (**Figures 2A and B**). The amino acid-substitution mutation R99E resulted in the production of anucleate cells regardless of the incubation temperature. However, filamentation at the restrictive temperature was remarkably suppressed by R99E, indicating an improvement in only temperature-sensitive growth, as observed in **Figure 1**.

**Figure 3.**
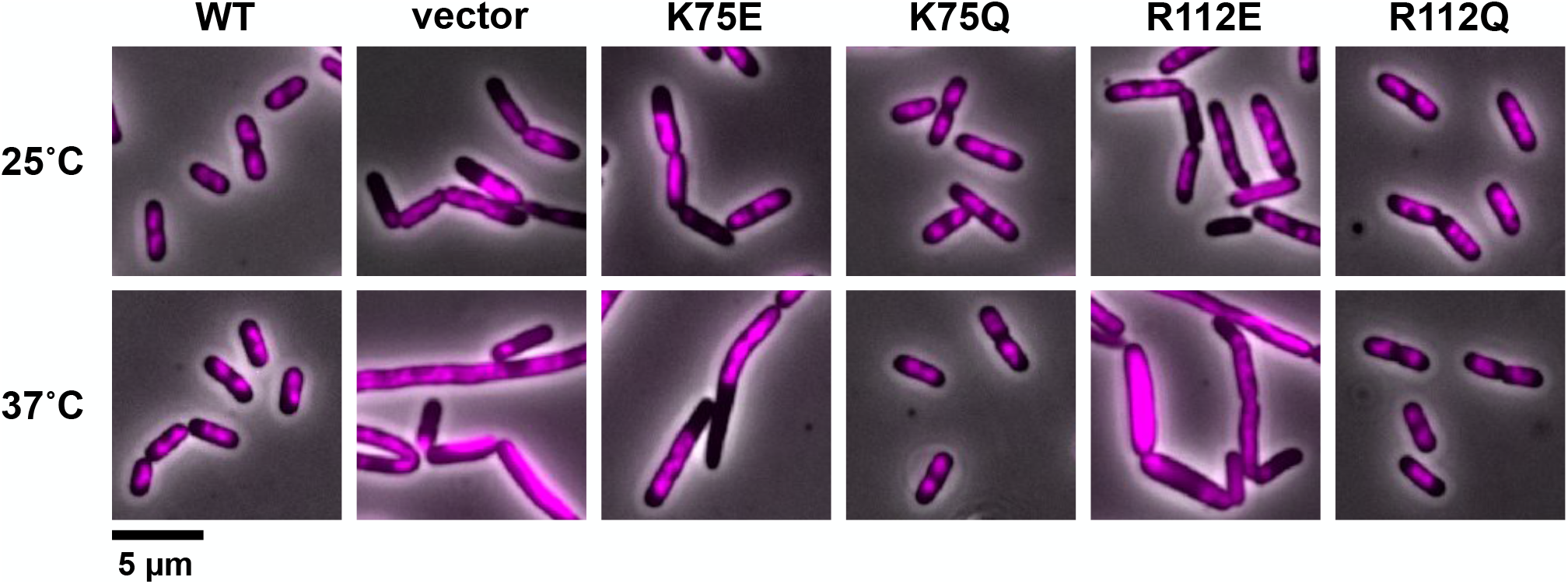
Nucleoid segregation in cells expressing the MukB head domain mutants. A series of YAN4081 (Δ*mukB*::*cat*) strains harboring the plasmid encoding the indicated His_6_-MukB mutant is depicted as merged images of phase-contrast images and DAPI-stained fluorescent images (pseudo color in magenta). The images of cells cultured at 25°C are displayed in the upper panels, and images at 37°C are shown in the lower panels. The scale bar indicates 5 µm.

### The DNA-binding activities of the amino acid substitution mutants

We next assessed the DNA-binding activities of the amino acid substitution mutants by means of an electrophoretic mobility shift assay (EMSA). The EMSA allows for the quantitative analysis of DNA-binding activities by measuring the delay in migration of DNA molecules caused by protein binding; purified MukB proteins binding to dsDNA and ssDNA. We quantified the intensity of DNA bands that did not bind to the purified MukB protein, and the DNA-binding activity was defined as the concentration of MukB protein required to shift half of the given amount of DNA, denoted as [MukB]_50_. The [MukB]_50_ values were calculated based on the binding kinetics of each mutated protein (**Figures 4, 5 and 6 and Table 2**). The wild-type His_6_-MukB^WT^ protein exhibited [MukB]_50_ values of 13.3 ± 4.1 for ssDNA and 12.0 ± 3.0 nM for dsDNA (**Figures 4, 5 and 6 and Table 2**).

**Figure 4.**
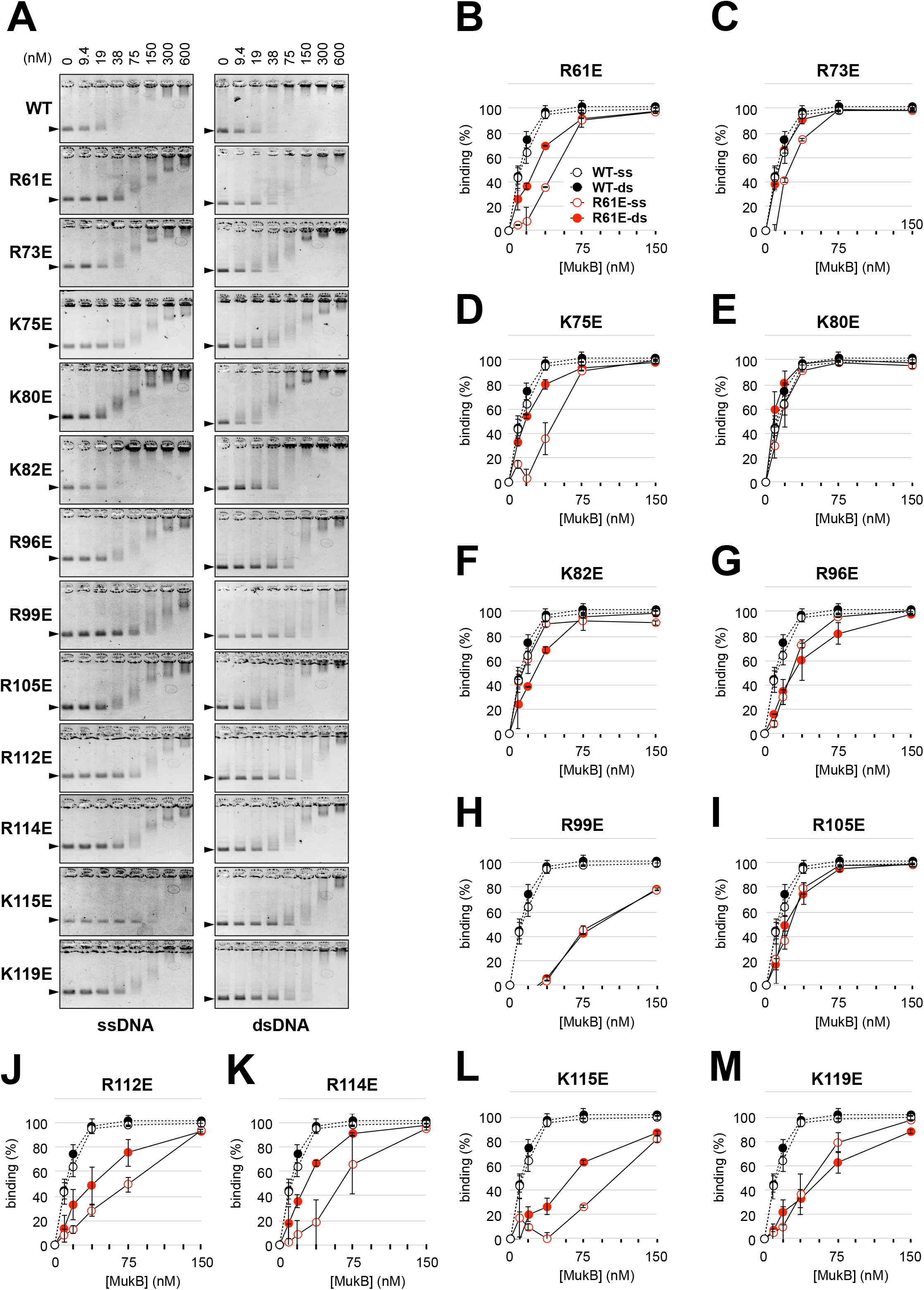
The ssDNA-binding and dsDNA-binding activities of the MukB head domain mutants with Glu (E) substitutions. (**A**) Gel images of the electromobility shift assay (EMSA). The amount of purified His_6_-MukB mutant proteins in each reaction is indicated in the top row. The left and the right panel show EMSA with cssDNA and cccDNA, respectively. Arrows indicate the positions of DNA to which the MukB protein does not bind. (**B**-**M**) Kinetics of the ssDNA-binding and dsDNA-binding activities of the MukB head domain mutants with Glu (E) substitutions were investigated. A single amino acid substitution of the MukB head domain mutants is indicated at the top of each graph. The intensities of the DNA bands to which the MukB protein does not bind in the EMSA gels in (**A**) are graphically shown according to inputs of the MukB protein, ranging from 0 to 150 nM. The mean values of the two or three independent experiments are indicated along with the standard deviations. The DNA-binding activities of the wild-type MukB protein are displayed for each reaction: black filled circles represent binding to dsDNA and black open circles represent binding to ssDNA. The DNA-binding activities of the mutated MukB protein are shown as follows: red filled circles represent binding to dsDNA and red open circles represent binding to ssDNA.

**Figure 5.**
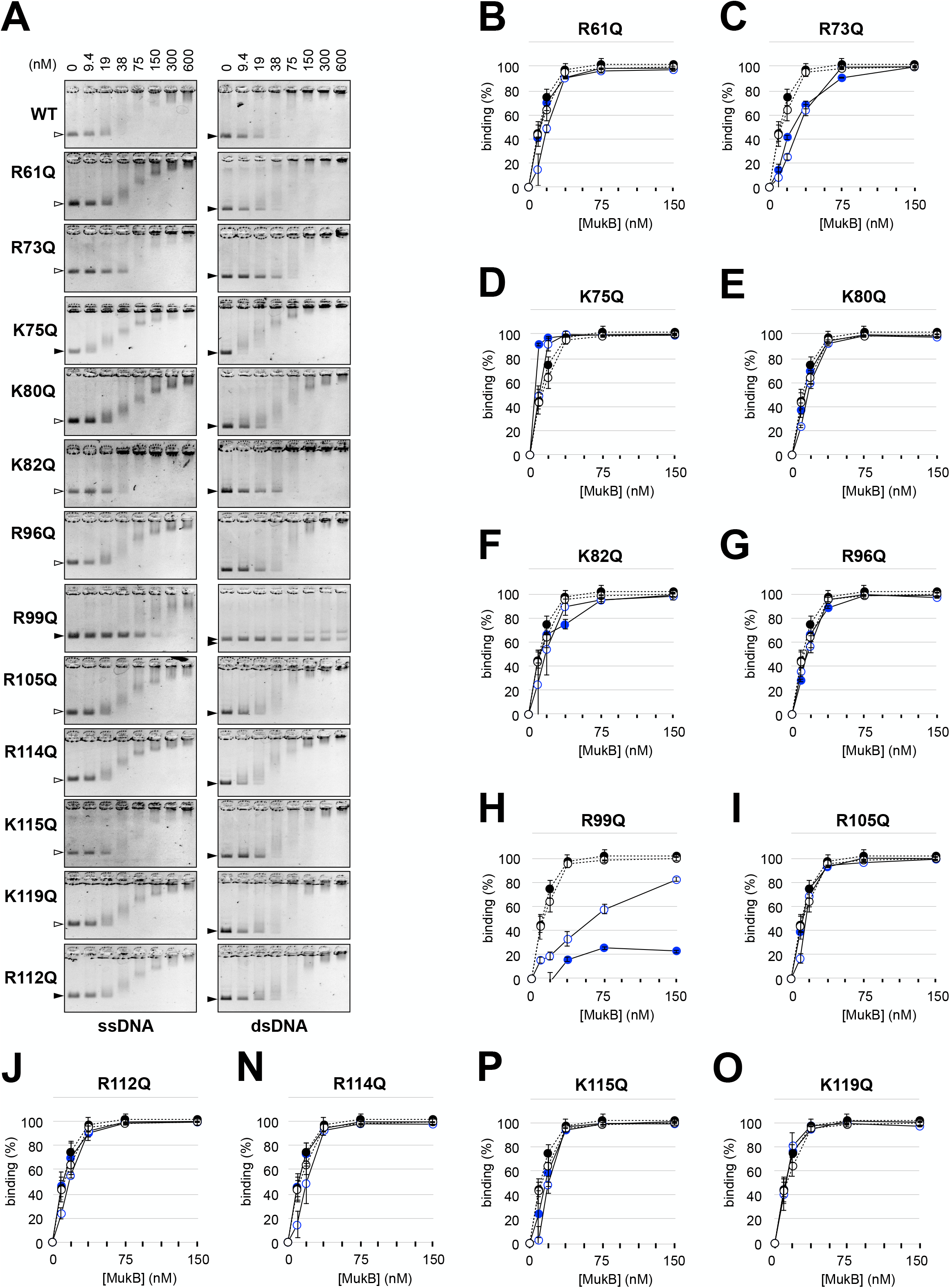
The ssDNA-binding and dsDNA-binding activities of the MukB head domain mutants with Gln (Q) substitutions. This figure is the same as Figure 4 except that the results are shown for Gln (Q) mutants instead of Glu (E) mutants.

**Figure 6.**
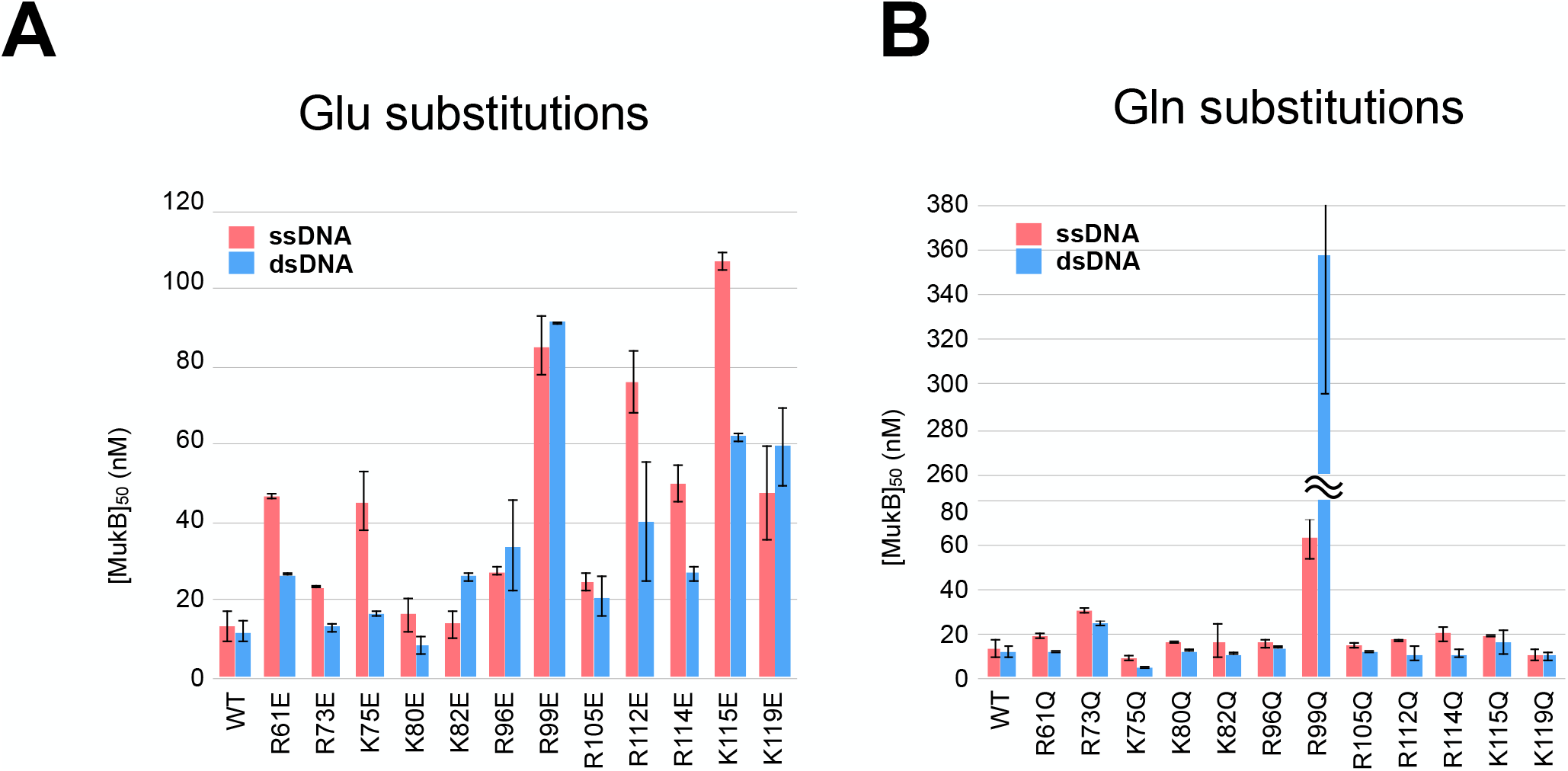
The [MukB]_50_ values as the ssDNA -binding and dsDNA-binding activities of the MukB head domain mutants. The [MukB]_50_ values are calculated based on the kinetics of the DNA-binding activities of the MukB head domain mutants in Figure 4 and 5. **The ssDNA-binding and dsDNA-binding** activities of the MukB head domain mutants, which are substituted to Glu (E), to ssDNA and dsDNA are indicated in panel A, and those of the Gln (Q)-substituted mutants are indicated in panel B. The mean values of the two or three independent experiments are indicated along with the standard deviations.

**Table 2.** The DNA-binding activities of the MukB head domain mutants to ssDNA and dsDNA. [MukB]_50_ values in two or three independent experiences are shown with the mean value ± standard deviations.

|  | binding to ssDNA |  |  |  | binding to dsDNA |  |  |  |
| --- | --- | --- | --- | --- | --- | --- | --- | --- |
| | 1st | 2nd | 3rd | mean ± SD | 1st | 2nd | 3rd | mean ± SD |
| WT | 14 | 18 | 8 | 13.3 ± 4.1 | 15 | 9 | | 12.0 ± 3.0 |
| R61E | 48 | 46 | | 47.0 ± 1.0 | 27 | 26 | | 26.5 ± 0.5 |
| R61Q | 21 | 18 | | 19.5 ± 1.5 | 12 | 13 | | 12.5 ± 0.5 |
| R73E | 23 | 24 | | 23.5 ± 0.5 | 14 | 12 | | 13.0 ± 1.0 |
| R73Q | 32 | 30 | | 31.0 ± 1.0 | 26 | 23 | | 24.5 ± 1.5 |
| K75E | 53 | 38 | | 45.5 ± 7.5 | 18 | 16 | 16 | 16.7 ± 0.9 |
| K75Q | 9 | 12 | 8 | 9.7 ± 1.7 | 5 | 6 | 5 | 5.3 ± 0.5 |
| K80E | 12 | 21 | | 16.5 ± 4.5 | 6 | 11 | | 8.5 ± 2.5 |
| K80Q | 16 | 16 | | 16.0 ± 0.0 | 12 | 14 | | 13.0 ± 1.0 |
| R96E | 29 | 26 | | 27.5 ± 1.5 | 46 | 22 | | 34.0 ± 12.0 |
| R96Q | 14 | 18 | | 16.0 ± 2.0 | 15 | 14 | | 14.5 ± 0.5 |
| R99E | 78 | 93 | | 85.5 ± 7.5 | 92 | 91 | | 91.5 ± 0.5 |
| R99Q | 71 | 54 | | 62.5 ± 8.5 | 420 | 295 | | 357.5 ± 62.5 |
| R105E | 22 | 27 | | 24.5 ± 2.5 | 26 | 16 | | 21.0 ± 5.0 |
| R105Q | 14 | 17 | | 15.5 ± 1.5 | 12 | 12 | | 12.0 ± 0.0 |
| R112E | 72 | 88 | 69 | 76.3 ± 8.3 | 19 | 45 | 57 | 40.3 ± 15.9 |
| R112Q | 17 | 18 | 18 | 17.7 ± 0.5 | 8 | 16 | 11 | 11.7 ± 3.3 |
| R114E | 55 | 45 | | 50.0 ± 5.0 | 26 | 30 | 25 | 27.0 ± 2.2 |
| R114Q | 22 | 24 | 15 | 20.3 ± 3.9 | 13 | 14 | 8 | 11.7 ± 2.6 |
| K115E | 110 | 105 | | 107.5 ± 2.5 | 63 | 61 | | 62.0 ± 1.0 |
| K115Q | 19 | 20 | | 19.5 ± 0.5 | 11 | 22 | | 16.5 ± 5.5 |
| K119E | 60 | 35 | | 47.5 ± 12.5 | 70 | 49 | | 59.5 ± 10.5 |
| K119Q | 8 | 14 | | 11.0 ± 3.0 | 13 | 8 | | 10.5 ± 2.5 |

The results for the Glu mutants revealed that the ssDNA-binding activities of the MukB proteins were decreased by the specific mutations; this was observed for each of the mutants, such as MukB**^R61E^**, MukB**^K75E^**, MukB**^R99E^**, MukB**^R112E^**, MukB**^R114E^**, MukB**^K115E^**, and MukB**^K119E^**. The [MukB]_50_ values for these mutants were more than two-fold higher compared to that of MukB**^WT^**. In the cases of MukB**^R99E^** and MukB**^K119E^**, the dsDNA-binding activities were also reduced to the same extent. Hence, the substitution of Glu for these amino acid residues equally affected both the ssDNA and dsDNA-binding abilities.

On the other hand, for the other mutants, the ssDNA-binding activities were markedly decreased compared to the dsDNA-binding activities. For example, the [MukB]_50_ value of MukB**^K75E^** for ssDNA was 45.5 ± 7.5 nM, whereas it was 16.7 ± 0.9 nM for dsDNA, which was comparable to the [MukB]_50_ value of MukB**^WT^** (12.0 ± 3.0 nM) (**Figures 4D, 5D and 6A**). In the case of MukB**^K75E^**, the ssDNA-binding activity was remarkably decreased, while the dsDNA-binding activity was not distinctly affected. Similar trends were observed for MukB**^R61E^** and MukB**^R114E^**, although their dsDNA-binding activities were slightly worse than that for MukB**^WT^** (**Figures 4B, 4K, 5B, 5K and 6A**). Additionally, MukB**^K112E^** and MukB**^K115E^** exhibited distinct differences in DNA-binding activities between ssDNA and dsDNA. Their ssDNA-binding activities were considerably lower than that of MukB**^WT^**, and the dsDNA-binding activities were also severely decreased (**Figures 4J, 4L, 5J, 5L and 6A**).

In contrast to the results observed with the Glu substitution mutants, the [MukB]_50_ values of the Gln substitution mutants were comparable to that of MukB**^WT^**, but not to those of MukB**^R73Q^** and MukB**^R99Q^** (**Figure 6B**). Notably, the dsDNA-binding activities of MukB**^R99Q^** were significantly reduced. The [MukB]_50_ value of MukB**^R99Q^** for dsDNA, 357.5 ± 62.5, was the highest among both the Glu and Gln substitution mutants (**Figures 5H and 6B**). Furthermore, the ssDNA-binding activity of MukB**^R99Q^**was also remarkably reduced, with a value of 62.5 ± 8.5 nM for ssDNA (**Figures 5H and 6B**).

### The topological DNA-binding activities of the amino acid substitution mutants

To assess the topological loading of MukB on DNA, we utilized the MU assay, which measures the amount of recovered DNA from a DNA-SMC-binding complex after eliminating the salt-sensitive binding of SMC protein to DNA {Niki.2016 }. In this assay, a circular single-stranded DNA substrate was incubated with purified histidine-tagged MukB under a low-salt condition (25 mM KCl). The DNA substrates bound to histidine-tagged MukB were then recovered using affinity beads that bind to the histidine-tag followed by washing with a high salt buffer solution (including 750 mM KCl). The recovered DNA substrates were analyzed by agarose gel electrophoresis, specifically by pulling down with histidine-tagged MukB (**Figure 7**). By employing this method, we were able to evaluate the topological loading activity of the mutated MukB protein.

**Figure 7.**
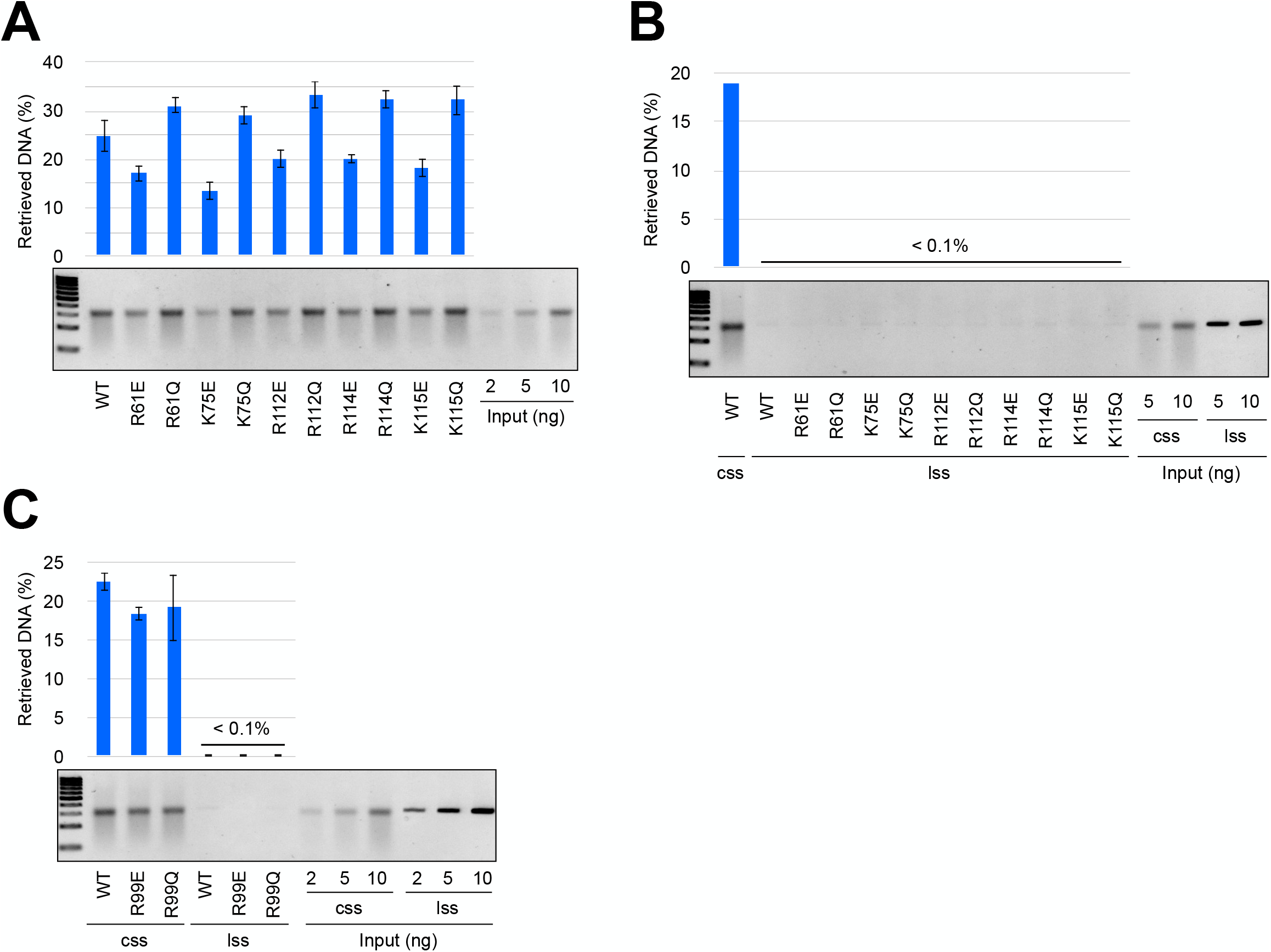
Topological DNA-binding activities of MukB mutants. (**A, C**) Circular single-stranded (css) DNA of pUC119 or (**B, C**) linear single-stranded (lss) DNA of pUC119. The mean of the two or three independent experiments are graphically shown with standard deviations.

We investigated whether the defects in the ssDNA-binding activities observed in the Glu substitutions (MukB**^R61E^**, MukB**^K75E^**, MukB**^R112E^**, MukB**^R114E^**, and MukB**^R115E^**) also affected their topological DNA-binding activities. Our experiments confirmed that all these Glu-substituted mutants exhibited decreased DNA recovery rates (**Figure 7A**), indicating that these mutations weakened the topological-binding activities. In contrast, the topological-binding activities of the Gln-substituted mutants remained equivalent to that of the WT protein, with MukB**^R112Q^** and MukB**^R114Q^** showing slightly decreased activities. When linear ssDNA was used instead of circular DNA, no DNA molecule were pulled down (**Figure 7B**). This result strongly suggests that the recovered cssDNA in this assay was topologically entrapped product. Collectively, the results indicate that the presence of a negative charge of the amino acid residues affects the topological-binding activity.

The mutants MukB**^R99E^** and MukB**^R99Q^** also affected the ssDNA-binding affinity in the EMSA assay. These substitutions differed from those of MukB**^R61E^**, MukB**^K75E^**, MukB**^R112E^**, MukB**^R114E^**, and MukB**^R115E^** in that they affected dsDNA-binding as much or more than they affected ssDNA-binding. Topological-binding activities of MukB**^R99E^** and MukB**^R99Q^** for the circular ssDNA were slightly lower than that of MukB**^WT^** but were maintained significantly (**Figure 7C**). The finding that no linear ssDNAs were retrieved showed that the DNAs were topologically entrapped (**Figure 7C**).

## DISCUSSION

Although some SMC proteins exhibit ssDNA-binding ability, the biological significance of their ssDNA-binding ability remains poorly understood. In this study, our objective was to identify the specific amino acid residues responsible for the ssDNA-binding ability of the MukB protein. We successfully identified several positively charged residues, namely R61, K75, R112, R114, and K115, located in the head domain of MukB, which play a major role in ssDNA-binding. Substituting Glu for these amino acid residues resulted in noticeable differences in the DNA-binding abilities between ssDNA and dsDNA. Notably, the K75E mutation alone reduced the ssDNA-binding ability while preserving the dsDNA-binding ability. The substitution of Glu for three of these crucial amino acid residues, R61, K75, and R112, significantly impacted all the phenotypes of cell growth, nucleoid separation, and topological loading.

In contrast, the Glu substitutions for the amino acid residues R114 and K115 did not affect cell growth or nucleoid separation. Nevertheless, the introduction of negative charge in these amino acid residues resulted in a reduction in the ssDNA-binding ability rather than the dsDNA-binding ability as well as a reduction in the topological loading ability onto ssDNA. The amino acids residues R114 and R115 locate at the basement of R112, and they support the protrusion of the charged residue R112 from the MukB head (**Figure 8**). Hence, it is reasonable to hypothesize that the amino acid residues R61, K75, and R112 collectively form the primary site for the ssDNA binding, while the two amino acid residues R114 and R115 assist in the efficient binding of ssDNA to the primary site.

**Figure 8.**
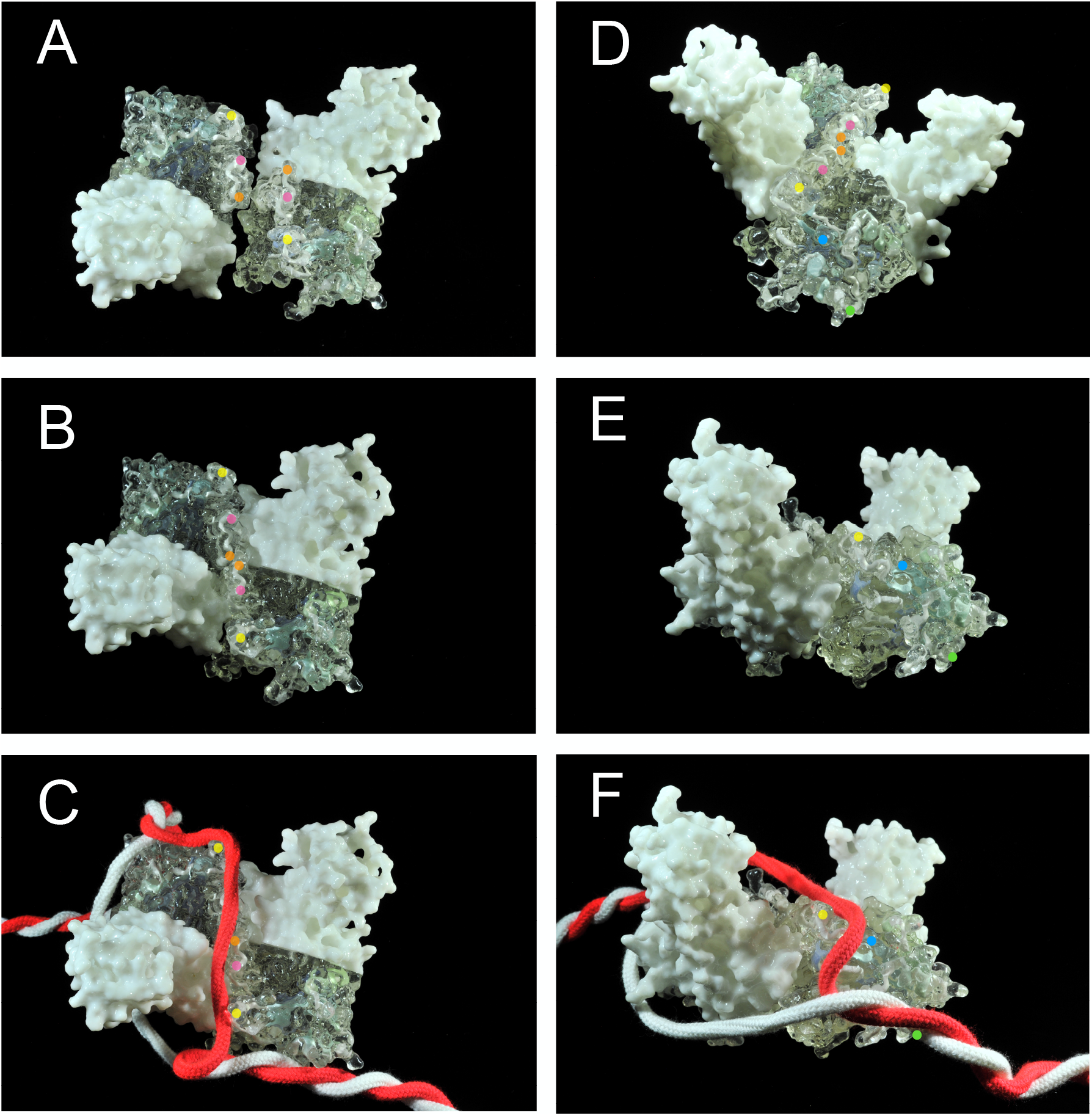
Model of the stable topological binding of MukB using the residues R61, K75 and R112. (**A**) A 3D model of the head domain of *Haemophilus ducreyi* MukB (PDB-ID: 3EUK) was made using a 3D printer (KawakamiModel; http://studio-midas.com/kawakamimodel/). In two monomers the 6 residues R61 (orange), K75 (magenta) and R112 (yellow) in each head are represented from the top view. (**B**) When the MukB dimer closes, these residues align within the inner surface of the closed MukB ring. (**C**) After MukB captures ssDNA inside its ring, ssDNA binds to the inner surface through the ssDNA-binding domain formed by aligned axis residues. The ssDNA crossing the MukB heads stabilizes the state of the engaged MukB head. The twisted strands (red and white) represent DNA with a melted part (**D**) The residues R99 (blue) and R119 (green) are represented along with the 6 residues R61 (orange), K75 (magenta) and R112 (yellow). (**E**) The residues R99 (blue) and R119 (green) are represented from the side view. (D) The residues R99 and R119 hold ssDNA or dsDNA.

The substitutions of the negatively charged Glu for R99 and K119 resulted in an equal reduction of binding ability to both dsDNA and ssDNA. Furthermore, substitution of the neutrally charged Gln for R99 led to the loss of these DNA-binding abilities, with the dsDNA-binding ability being the least weakened among all the amino acid substitutions in this experiment. However, despite the decrease in DNA-binding abilities, the Glu and Gln substitutions for R99 and K119 never showed defects in cell growth at high temperatures. Thus, the DNA-binding abilities of R99 and K119 are not essential for the basic biological function of MukB. Taken together, these results suggest the likelihood that there are at least two DNA-binding domains within the MukB head. One involves the alignment of R61, K75, and R112 for an essential ssDNA binding. The other domain consists of R99 and K119, which are involved in a non-essential DNA binding to ssDNA and dsDNA, especially to dsDNA. This non-essential DNA binding to dsDNA could potentially facilitate to maintain melted DNA onto the MukB dimer as shown below. Alternatively, it could contribute to the distribution of MukB to the whole chromosome region except the *ter* domain.

As to why the MukB dimer exhibits a preference for entrapping ssDNA over dsDNA, we propose the following scenario based on the spatial position of the above-described residues within the protein structure model of the MukB head. The in vitro assay showed that a purified MukB dimer repeatedly opens and closes its ring structure through thermal motion rather than ATP hydrolysis (39. The residues R61, K75 and R112 are positioned on the inner surface of the MukB head (**Figure 8A**). When the head domains of the MukB dimer bind with each other and form a closed form of the head, these six amino acid residues align within the inner surface of the closed MukB head (**Figure 8B**).

Consequently, it is plausible that the alignment of R61, K75, and R112 forms an axis of positively charged amino acid residues, functioning as a binding site almost exclusively for ssDNA only when the MukB ring is closed. Consider the case in which the axis amino acid residues bind to entrapped ssDNA within the ring of the MukB dimer. In that case, the ssDNA can span across the closed dimer head, thereby stabilizing the closed form while the ssDNA interacts with the axis amino acid residues (**Figure 8C**). Furthermore, R99, which is located diagonally below R112, may assist in additionally binding the extended ssDNA in the appropriate orientation against the MukB head (**Figure 8D**). Overall, in the ssDNA-bound MukB dimer, it appears that the heads of the dimer are tied by the ssDNA; consequently, the ssDNA acts as a linking element with each head of the MukB dimer (**Figure 8E**).

The substitutions R61Q, K75Q, and R112Q maintained the DNA-binding activities and topological loading activities at the wild-type level. However, the R61Q substitution alone caused defects in cell growth and nucleoid segregation. The structural information suggests that the amino acid residue equivalent to R61 is directly involved in the hydrolysis of ATP of the ATP-binding cassette transporters (19, 60). The R61 amino acid residue corresponds to the residue of the Arg finger, which is the motif involved in ATPase hydrolysis (60. In the case of the R61Q substitution, it is conceivable that the residue is involved not only in DNA-binding ability but also in the ATPase activity, which is necessary for the complete biological function of MukB. Furthermore, the K80 amino acid residue is also involved in the ATPase activity since it is positioned in such a way that its side chain faces an ATP molecule, as indicated by the crystal structure of the ATP-binding cassette transporter (19. Therefore, it is reasonable that both the K80E and K80Q substitutions caused defects in cell growth and nucleoid segregation, although they did not affect the DNA-binding activities.

The temperature-dependent cell growth is a common characteristic observed in both the *mukB*-deletion mutant and the *smc*-deletion mutant of *B. subtilis*. In both cases, the deletion mutants produced filamentous cells and exhibited errors in chromosome segregation at high temperatures. However, the mechanisms that could explain why these bacterial condensin mutants displayed temperature-dependent cell growth and why this defect became worse in the nutrient-rich medium remain elusive. It is believed that the progression of the multiple replication forks, which was enhanced under the nutrient-rich medium condition and under higher growth temperature, contributes to the disruption of nucleoid compaction in the absence of bacterial condensin. Interestingly, the R99E amino acid substitution mutant was found to suppress the temperature-dependent cell growth observed in the *mukB*-deletion mutant. However, this mutant still produced filamentous cells and exhibited errors in chromosome segregation at a high temperature. Therefore, cell elongation was not the sole cause of the temperature-dependent cell growth. It appears that the R99 amino acid residue is involved in a latent and unsuspected function of MukB that contributed to this phenomenon.

The *B. subtilis* Smc-ScpAB complex has been observed to accumulate on the actively transcribed ribosomal RNA (*rrn*) operon (40. Within this operon, an ssDNA segment that facilitates the accumulation of Smc-ScpAB has been identified. (53. Hence, it is plausible that preferential topological loading onto ssDNA could be utilized to promote the accumulation of bacterial condensin in specific regions such as the *rrn* operon. This is consistent with previous ChiP analysis by Gruber and Errington (61 that showed the enrichment of the Bs Smc on *rrn* regions. Another ChiP analysis showed that yeast condensin was accumulated at the highly transcribed regions including rDNA (62. The ssDNA-targeted loading is a common feature for both the prokaryotic and eukaryotic condensins.

The ssDNA-binding of MukB in this report is the first step in the functional cycle of bacterial condensin. The behavior of the MukB after being loaded onto the ssDNA region should be investigated to better understand the mechanism of nucleoid compaction. To understand the whole picture of this cycle, the roles of the MukEF, the non-SMC subunits, and the hydrolysis of ATP must be further investigated.

## DATA AVAILABILITY

There is no relevant data availability.

## SUPPLEMENTARY DATA

The Supplementary Data are available online.

## ACKNOWLEDGEMENTS

We thank all members of the Niki Laboratory for their technical support. We thank the National Bioresource Project (NBRP) for providing materials. KA thanks the National Institute of Genetics for providing a position as part of the NIG Postdoctoral Fellowship.

## FUNDING

This work was supported by JJSPS KAKENHI Grants to KA (no. JP17H07328) and HN (nos. JP18H02485 and JP23H02525).

## CONFLICT OF INTEREST

None of the authors has any conflict of interest to report.

---

**Supplementary figure 1. Nucleoid segregation in cells expressing head domain mutants.**

In addition to the strains shown in Figure 3, a series of YAN4081 (Δ*mukB*::*cat*) strains harboring the plasmid encoding the indicated His_6_-MukB mutant is depicted as merged images of phase-contrast images and DAPI-stained fluorescent images (pseudo color in magenta). The images of cells cultured at 25°C are displayed in the upper panels, and images at 37°C are shown in the lower panels. The scale bar indicates 5 µm.

**Supplementary figure 2. Histogram of frequencies of the anucleate cell production.**

Frequencies of anucleate cell production in Table 1 are represented as a histogram.

## Notes

### Competing Interest Statement

The authors have declared no competing interest.

## REFERENCES

1. Kato, J., Nishimura, Y., Imamura, R., Niki, H., Hiraga, S. and Suzuki, H. (1990) New topoisomerase essential for chromosome segregation in E. coli. Cell, 63, 393–404.

2. Niki, H., Jaffé, A., Imamura, R., Ogura, T. and Hiraga, S. (1991) The new gene mukB codes for a 177 kd protein with coiled-coil domains involved in chromosome partitioning of E. coli. EMBO J., 10, 183–193.

3. Sawitzke, J.A. and Austin, S. (2000) Suppression of chromosome segregation defects of Escherichia coli muk mutants by mutations in topoisomerase I. Proc. Natl. Acad. Sci., 97, 1671–1676.

4. Cui, Y., Petrushenko, Z.M. and Rybenkov, V.V. (2008) MukB acts as a macromolecular clamp in DNA condensation. Nat. Struct. Mol. Biol., 15, 411–418.

5. Nasmyth, K. and Haering, C.H. (2005) THE STRUCTURE AND FUNCTION OF SMC AND KLEISIN COMPLEXES. Annu. Rev. Biochem., 74, 595–648.

6. Nolivos, S. and Sherratt, D. (2014) The bacterial chromosome: architecture and action of bacterial SMC and SMC-like complexes. FEMS Microbiol. Rev., 38, 380–392.

7. Haering, C.H. and Gruber, S. (2016) SnapShot: SMC Protein Complexes Part I. Cell, 164, 326–326.e1.

8. Hirano, T. (2016) Condensin-Based Chromosome Organization from Bacteria to Vertebrates. Cell, 164, 847–857.

9. Mäkelä, J. and Sherratt, D. (2020) SMC complexes organize the bacterial chromosome by lengthwise compaction. Curr Genet, 338, 528–5.

10. Britton, R.A., Lin, D.C.-H. and Grossman, A.D. (1998) Characterization of a prokaryotic SMC protein involved in chromosome partitioning. Genes Dev., 12, 1254–1259.

11. Moriya, S., Tsujikawa, E., Hassan, A.K.M., Asai, K., Kodama, T. and Ogasawara, N. (1998) A Bacillus subtilis gene-encoding protein homologous to eukaryotic SMC motor protein is necessary for chromosome partition. Mol. Microbiol., 29, 179–187.

12. Mascarenhas, J., Soppa, J., Strunnikov, A.V. and Graumann, P.L. (2002) Cell cycle-dependent localization of two novel prokaryotic chromosome segregation and condensation proteins in Bacillus subtilis that interact with SMC protein. EMBO J., 21, 3108–3118.

13. Yamanaka, K., Ogura, T., Niki, H. and Hiraga, S. (1996) Identification of two new genes, mukE andmukF, involved in chromosome partitioning inEscherichia coli. Mol. Gen. Genet., 250, 241– 251.

14. Niki, H., Imamura, R., Kitaoka, M., Yamanaka, K., Ogura, T. and Hiraga, S. (1992) E.coli MukB protein involved in chromosome partition forms a homodimer with a rod-and-hinge structure having DNA binding and ATP/GTP binding activities. EMBO J., 11, 5101–5109.

15. Matoba, K., Yamazoe, M., Mayanagi, K., Morikawa, K. and Hiraga, S. (2005) Comparison of MukB homodimer versus MukBEF complex molecular architectures by electron microscopy reveals a higher-order multimerization. Biochem. Biophys. Res. Commun., 333, 694–702.

16. Bürmann, F., Lee, B.-G., Than, T., Sinn, L., O’Reilly, F.J., Yatskevich, S., Rappsilber, J., Hu, B., Nasmyth, K. and Löwe, J. (2019) A folded conformation of MukBEF and cohesin. Nat. Struct. Mol. Biol., 26, 227–236.

17. Yamazoe, M., Onogi, T., Sunako, Y., Niki, H., Yamanaka, K., Ichimura, T. and Hiraga, S. (1999) Complex formation of MukB, MukE and MukF proteins involved in chromosome partitioning in Escherichia coli. EMBO J., 18, 5873–5884.

18. Petrushenko, Z.M., Lai, C.-H. and Rybenkov, V.V. (2006) Antagonistic Interactions of Kleisins and DNA with Bacterial Condensin MukB*. J. Biol. Chem., 281, 34208–34217.

19. Woo, J.-S., Lim, J.-H., Shin, H.-C., Suh, M.-K., Ku, B., Lee, K.-H., Joo, K., Robinson, H., Lee, J., Park, S.-Y., et al. (2009) Structural Studies of a Bacterial Condensin Complex Reveal ATP-Dependent Disruption of Intersubunit Interactions. Cell, 136, 85–96.

20. Palecek, J.J. and Gruber, S. (2015) Kite Proteins: a Superfamily of SMC/Kleisin Partners Conserved Across Bacteria, Archaea, and Eukaryotes. Structure, 23, 2183–2190.

21. Kamada, K., Su’etsugu, M., Takada, H., Miyata, M. and Hirano, T. (2017) Overall Shapes of the SMC-ScpAB Complex Are Determined by Balance between Constraint and Relaxation of Its Structural Parts. Structure, 25, 603–616.e4.

22. Zawadzka, K., Zawadzki, P., Baker, R., Rajasekar, K.V., Wagner, F., Sherratt, D.J. and Arciszewska, L.K. (2018) MukB ATPases are regulated independently by the N-and C-terminal domains of MukF kleisin. eLife, 7, e31522.

23. Rajasekar, K.V., Baker, R., Fisher, G.L.M., Bolla, J.R., Mäkelä, J., Tang, M., Zawadzka, K., Koczy, O., Wagner, F., Robinson, C.V., et al. (2019) Dynamic architecture of the Escherichia coli structural maintenance of chromosomes (SMC) complex, MukBEF. Nucleic Acids Res., 47, 9696–9707.

24. Vos, S.M., Stewart, N.K., Oakley, M.G. and Berger, J.M. (2013) Structural basis for the MukB-topoisomerase IV interaction and its functional implications in vivo. EMBO J., 32, 2950–2962.

25. Nicolas, E., Upton, A.L., Uphoff, S., Henry, O., Badrinarayanan, A. and Sherratt, D. (2014) The SMC Complex MukBEF Recruits Topoisomerase IV to the Origin of Replication Region in Live Escherichia coli. mBio, 5, e01001–13.

26. Kumar, R., Nurse, P., Bahng, S., Lee, C.M. and Marians, K.J. (2017) The MukB–topoisomerase IV interaction is required for proper chromosome compaction. J. Biol. Chem., 292, 16921–16932.

27. Bürmann, F., Funke, L.F.H., Chin, J.W. and Löwe, J. (2021) Cryo-EM structure of MukBEF reveals DNA loop entrapment at chromosomal unloading sites. Mol. Cell, 81, 4891–4906.e8.

28. Prince, J.P., Bolla, J.R., Fisher, G.L.M., Mäkelä, J., Fournier, M., Robinson, C.V., Arciszewska, L.K. and Sherratt, D.J. (2021) Acyl carrier protein promotes MukBEF action in Escherichia coli chromosome organization-segregation. Nat. Commun., 12, 6721.

29. Bahng, S., Kumar, R. and Marians, K.J. (2022) Intersubunit and intrasubunit interactions driving the MukBEF ATPase. J. Biol. Chem., 298, 101964.

30. Gloyd, M., Ghirlando, R. and Guarné, A. (2011) The Role of MukE in Assembling a Functional MukBEF Complex. J. Mol. Biol., 412, 578–590.

31. Hirano, M. and Hirano, T. (2006) Opening Closed Arms: Long-Distance Activation of SMC ATPase by Hinge-DNA Interactions. Mol. Cell, 21, 175–186.

32. Bürmann, F., Basfeld, A., Nunez, R.V., Diebold-Durand, M.-L., Wilhelm, L. and Gruber, S. (2017) Tuned SMC Arms Drive Chromosomal Loading of Prokaryotic Condensin. Mol. Cell, 65, 861–872.e9.

33. Li, Y., Schoeffler, A.J., Berger, J.M. and Oakley, M.G. (2010) The Crystal Structure of the Hinge Domain of the Escherichia coli Structural Maintenance of Chromosomes Protein MukB. J. Mol. Biol., 395, 11–19.

34. Ku, B., Lim, J., Shin, H., Shin, S. and Oh, B. (2010) Crystal structure of the MukB hinge domain with coiled-coil stretches and its functional implications. Proteins, 78, 1483–1490.

35. Murayama, Y. and Uhlmann, F. (2014) Biochemical reconstitution of topological DNA binding by the cohesin ring. Nature, 505, 367–371.

36. Uhlmann, F. (2016) SMC complexes: from DNA to chromosomes. Nat. Rev. Mol. Cell Biol., 17, 399–412.

37. Higashi, T.L. and Uhlmann, F. (2022) SMC complexes: Lifting the lid on loop extrusion. Curr. Opin. Cell Biol., 74, 13–22.

38. Wilhelm, L., Bürmann, F., Minnen, A., Shin, H.-C., Toseland, C.P., Oh, B.-H. and Gruber, S. (2015) SMC condensin entraps chromosomal DNA by an ATP hydrolysis dependent loading mechanism in Bacillus subtilis. eLife, 4, e06659.

39. Niki, H. and Yano, K. (2016) In vitro topological loading of bacterial condensin MukB on DNA, preferentially single-stranded DNA rather than double-stranded DNA. Sci. Rep., 6, 29469.

40. Yano, K. and Niki, H. (2017) Multiple cis-Acting rDNAs Contribute to Nucleoid Separation and Recruit the Bacterial Condensin Smc-ScpAB. Cell Rep., 21, 1347–1360.

41. Keyamura, K. and Hishida, T. (2019) Topological DNA-binding of structural maintenance of chromosomes-like RecN promotes DNA double-strand break repair in Escherichia coli. *Commun*. Biol., 2, 413.

42. Nasmyth, K. (2001) DISSEMINATING THE GENOME: Joining, Resolving, and Separating Sister Chromatids During Mitosis and Meiosis. Annu. Rev. Genet., 35, 673–745.

43. Alipour, E. and Marko, J.F. (2012) Self-organization of domain structures by DNA-loop-extruding enzymes. Nucleic Acids Res., 40, 11202–11212.

44. Goloborodko, A., Imakaev, M.V., Marko, J.F. and Mirny, L. (2016) Compaction and segregation of sister chromatids via active loop extrusion. eLife, 5, e14864.

45. Wang, X., Brandão, H.B., Le, T.B.K., Laub, M.T. and Rudner, D.Z. (2017) Bacillus subtilis SMC complexes juxtapose chromosome arms as they travel from origin to terminus. Science, 355, 524–527.

46. Ganji, M., Shaltiel, I.A., Bisht, S., Kim, E., Kalichava, A., Haering, C.H. and Dekker, C. (2018) Real-time imaging of DNA loop extrusion by condensin. Science, 360, 102–105.

47. Marko, J.F., De Los Rios, P., Barducci, A. and Gruber, S. (2019) DNA-segment-capture model for loop extrusion by structural maintenance of chromosome (SMC) protein complexes. Nucleic Acids Res., 47, 6956–6972.

48. Gerguri, T., Fu, X., Kakui, Y., Khatri, B.S., Barrington, C., Bates, P.A. and Uhlmann, F. (2021) Comparison of loop extrusion and diffusion capture as mitotic chromosome formation pathways in fission yeast. Nucleic Acids Res., 49, 1294–1312.

49. Lioy, V.S., Cournac, A., Marbouty, M., Duigou, S., Mozziconacci, J., Espéli, O., Boccard, F. and Koszul, R. (2018) Multiscale Structuring of the E. coli Chromosome by Nucleoid-Associated and Condensin Proteins. Cell, 172, 771–783.e18.

50. Mäkelä, J. and Sherratt, D.J. (2020) Organization of the Escherichia coli Chromosome by a MukBEF Axial Core. Mol. Cell, 78, 250–260.e5.

51. Danilova, O., Reyes-Lamothe, R., Pinskaya, M., Sherratt, D. and Possoz, C. (2007) MukB colocalizes with the oriC region and is required for organization of the two Escherichia coli chromosome arms into separate cell halves. Mol. Microbiol., 65, 1485–1492.

52. Badrinarayanan, A., Reyes-Lamothe, R., Uphoff, S., Leake, M.C. and Sherratt, D.J. (2012) In Vivo Architecture and Action of Bacterial Structural Maintenance of Chromosome Proteins. Science, 338, 528–531.

53. Yano, K., Noguchi, H. and Niki, H. (2022) Profiling a single-stranded DNA region within an rDNA segment that affects the loading of bacterial condensin. iScience, 25, 105504.

54. Sutani, T. and Yanagida, M. (1997) DNA renaturation activity of the SMC complex implicated in chromosome condensation. Nature, 388, 798–801.

55. Cuylen, S., Metz, J. and Haering, C.H. (2011) Condensin structures chromosomal DNA through topological links. Nat. Struct. Mol. Biol., 18, 894–901.

56. Johzuka, K. and Horiuchi, T. (2009) The cis Element and Factors Required for Condensin Recruitment to Chromosomes. Mol. Cell, 34, 26–35.

57. Murayama, Y., Samora, C.P., Kurokawa, Y., Iwasaki, H. and Uhlmann, F. (2018) Establishment of DNA-DNA Interactions by the Cohesin Ring. Cell, 172, 465–477.e15.

58. Datsenko, K.A. and Wanner, B.L. (2000) One-step inactivation of chromosomal genes in Escherichia coli K-12 using PCR products. Proc. Natl. Acad. Sci., 97, 6640–6645.

59. Nozaki, S. and Niki, H. (2019) Exonuclease III (XthA) Enforces In Vivo DNA Cloning of Escherichia coli To Create Cohesive Ends. J. Bacteriol., 201, e00660–18.

60. Lammens, A., Schele, A. and Hopfner, K.-P. (2004) Structural Biochemistry of ATP-Driven Dimerization and DNA-Stimulated Activation of SMC ATPases. Curr. Biol., 14, 1778–1782.

61. Gruber, S. and Errington, J. (2009) Recruitment of Condensin to Replication Origin Regions by ParB/SpoOJ Promotes Chromosome Segregation in B. subtilis. Cell, 137, 685–696.

62. Sutani, T., Sakata, T., Nakato, R., Masuda, K., Ishibashi, M., Yamashita, D., Suzuki, Y., Hirano, T., Bando, M. and Shirahige, K. (2015) Condensin targets and reduces unwound DNA structures associated with transcription in mitotic chromosome condensation. Nat. Commun., 6, 7815.

