## Supplemental Figures 1 and 2 for "Amino acid residues for specific binding to ssDNA facilitate topological loading of bacterial condensin MukB"

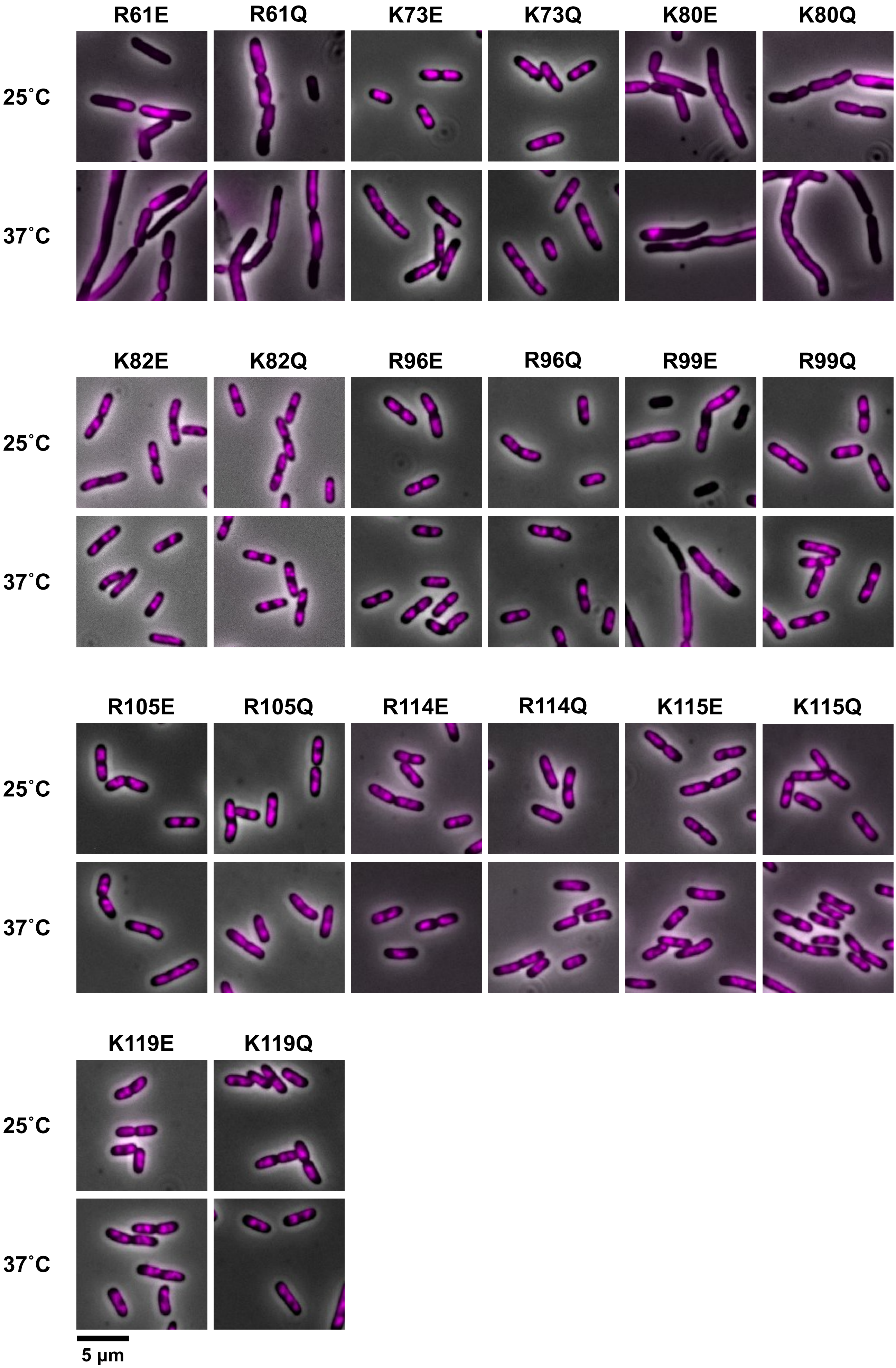


**Supplementary figure 1**

**Supplementary figure 1. Nucleoid segregation in cells expressing head domain mutants.**

In addition to the strains shown in Figure 3, a series of YAN4081 (D*mukB*::*cat*) strains harboring the plasmid encoding the indicated $His_6$-MukB mutant is depicted as merged images of phase-contrast images and DAPI-stained fluorescent images (pseudo color in magenta). The images of cells cultured at 25°C are displayed in the upper panels, and images at 37°C are shown in the lower panels. The scale bar indicates 5 μm.

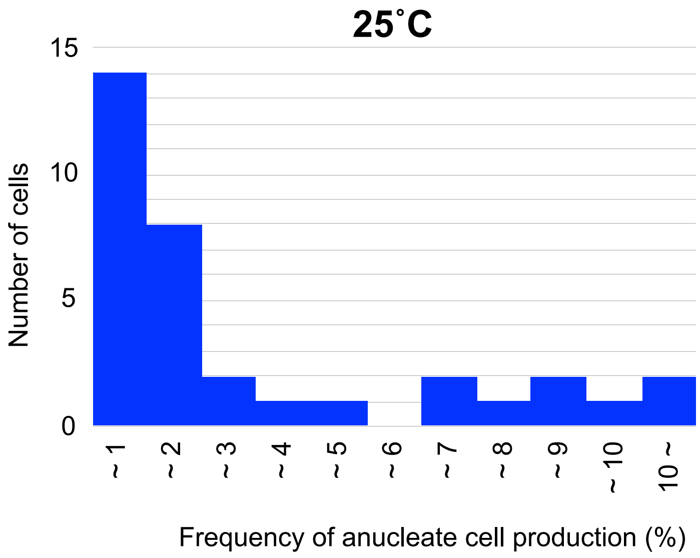


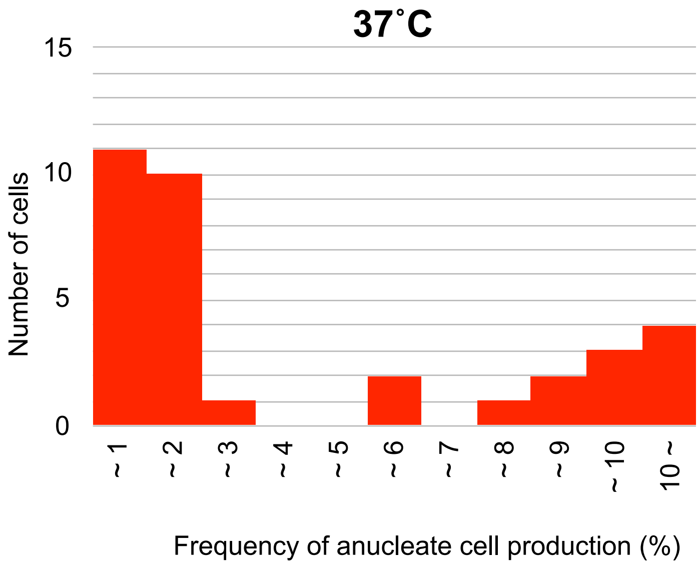


**Supplementary figure 2. Histogram of frequencies of the anucleate cell production.**
Frequencies of anucleate cell production in Table 1 are represented as a histogram.

### Supplementary Figure 2
